# Dynamic alterations in gene co-expression networks and gene-transcript associations characterize co-morbidities in cocaine use disorder

**DOI:** 10.1101/2024.07.10.602908

**Authors:** Chinwe Nwaneshiudu, Kiran Girdhar, Steven P. Kleopoulos, John F. Fullard, Eduardo R. Butelman, Muhammad A. Parvaz, Rita Z. Goldstein, Nelly Alia-Klein, Panos Roussos

## Abstract

**Background:** Individuals with cocaine use disorder (CUD) who attempt abstinence experience craving and relapse, which poses challenges in treatment. Longitudinal studies linking behavioral manifestations in CUD to the blood transcriptome in living individuals are limited. Therefore, we investigated the connection between drug use behaviors during abstinence with blood transcriptomics.

**Methods:** We conducted a comprehensive longitudinal study involving 12 subjects (9 males, 3 females) with CUD and RNA sequencing on blood collected at a drug-free baseline, and 3, 6 & 9 months thereafter. We categorized subjects into 2 responder groups (high-low) based on scores of drug use variables, and 3 responder groups (low-intermediate-high) on days of abstinence. We investigated differential expression and gene-transcript associations across responder groups at each time point. Lastly, we examined genes that are both co-expressed and showed dynamic expression with time.

**Results:** Genes with significant transcript associations between high and. intermediate days of abstinence at 9 months were notably enriched for cannabis use disorder, drinks weekly, and coronary artery disease risk genes. Time-specific gene co-expression analysis prioritized transcripts related to immune processes, cell cycle, RNA-protein synthesis, and second messenger signaling for days of abstinence.

**Conclusion:** We demonstrate that abstinence reflects robust changes in drug use behaviors and the blood transcriptome in CUD. We also highlight the importance of longitudinal studies to capture complex biological processes during abstinence in CUD.

## Introduction

Craving and enhanced reactivity to drug cues are core symptoms of drug addiction and are attributed to impairments in self-control with drug intake. Cocaine is one of the most commonly used illicit psychostimulants in the US and worldwide [1]. However, only 16% of individuals who use cocaine sporadically will progress to cocaine use disorder (CUD) [2], characterized by cycles of intoxication, withdrawal, and craving, with cardiovascular morbidity and other health, economic, social, and legal consequences [3] including mortality. CUD is also diagnosed based on the severity of symptoms (mild, moderate, and severe), which describe impaired control in the context of drug intake, risk-taking behaviors, social dysfunction as well as pharmacological effects with tolerance and withdrawal [4]. Cocaine use is also associated with accelerated aging as well as with acute and chronic vascular consequences causing cerebrovascular and cardiovascular diseases [5–7]. In contrast to other types of substance use disorders, there are no FDA-approved treatments for CUD [8]. Thus, a comprehensive understanding of the time course of cocaine use, abstinence, and relapse in individuals with CUD is essential to develop new treatments in future studies.

Treatment-seeking individuals who abstain from problematic cocaine use undergo a difficult course of unsuccessful recovery. Over 30.4% of patients eventually drop out and relapse back into regular cocaine and other drug use [9] and higher drop-out rates are observed in moderate and heavy users (51.8-66.7%), pointing to increased difficulties in maintaining abstinence with more severe cocaine use. Indeed, during attempted abstinence from cocaine use, lapses back to drug use are frequent and are preceded by craving - an intense experience of wanting to take the drug [10–12]. Craving is dynamic, time-dependent and is self-reported in response to either drug related images or other cues [13–15], or is generalized, unprovoked by images [16]. These persistent cravings, cue-craving and generalized craving, along with stress and other withdrawal symptoms [17] are good predictors of relapse to cocaine use [18] even when individuals undergo abstinence during clinical treatment [19]. By tracking the molecular underpinnings during abstinence, we can differentiate individuals with a relapsing pattern from those who successfully abstain over time, through their molecular dynamics.

Substance use disorders including CUD and other relevant traits are polygenic, heterogeneous, and multifactorial [20]. Most genetic and genomic studies to date are conducted in post-mortem brains, thus missing the crucial examination of dynamic behavioral, neurobiological, and environmental influences in gene expression associated with cocaine use [21]. The application of genome-wide transcriptomics to study CUD in living individuals is currently limited by the few available analytical techniques and access to relevant biological specimens, especially repeated specimens from the same individual longitudinally that are feasible only in non-invasive tissue collection. However, studies in this fashion are lacking. One study in women undergoing inpatient treatment for CUD was reported to show a positive association of microRNA (serum miR-181) expression in blood with higher dependence levels, and higher withdrawal symptoms but not for relapse during abstinence [22]. Another study examined the longitudinal association between objective changes in cocaine consumption and psychiatric symptoms in a mixed cohort of individuals (control, increased use, chronic cocaine use) at baseline and after a 1-year interval, which demonstrated that individuals who decreased cocaine consumption from a baseline visit till after 1-year had reduced mRNA expression of the glucocorticoid receptor gene NR3C1 in peripheral blood that approached similar mRNA levels to controls [23]. The use of both clinical and molecular biomarkers to identify vulnerabilities toward cocaine relapse or promote abstinence remains a focus of much interest [24, 25]. Longitudinal study designs with whole genome wide transcriptomics are well suited in this fashion, however, they are also computationally complex. In this study, we present an approach to examine dynamic changes in gene expression over time that occur with abstinence in CUD. This approach can further inform treatment strategies to promote longer abstinence and predict the propensity to relapse in vulnerable individuals.

Our analytical pipeline for longitudinal genome-wide gene expression was applied in an outpatient cohort of individuals with CUD. We show novel applications of blood-based transcriptomics as a potentially accessible and practical biomarker to examine gene expression variability over time during attempted abstinence in treatment-seeking individuals with CUD.

## Results

### Longitudinal study design and harmonization of drug use variables

We conducted a longitudinal study involving 12 abstinence-seeking subjects (9 males, 3 females) with CUD at regular intervals including a drug-free baseline, and at 3, 6 and 9 months time points. Over the course of the study, we gathered peripheral blood samples and the following drug use variables: cocaine cue-craving, generalized craving, cocaine withdrawal symptoms, perceived stress scores, and days of abstinence at each of the time points (see **Table 1** and **Fig. 1A**).

**Table 1.** Study demographics and drug use history of subjects at baseline visit.

|  |  |
| --- | --- |
| Sex (M/F) | 9/3 |
| Age (years) | 53.1(5.3) |
| Race (Caucasian/African American/Other) | 1/11/0 |
| Education (years in school) | 12.8 ± 1.8 |
| Depression (BDI) | 11.9 ± 10.5 |
| Anxiety (STAI total) | 76.9 ± 23.0 |
| Age of onset cocaine use (years) | 24.4 ± 27.3 |
| Age of regular cocaine use (years) | 27.1 ± 26.3 |
| Years of regular cocaine use (years) | 18.6 ± 9.3 |
| Frequency of yearly cocaine use (days per year) | 60 (19.5-162)* |
| Grams of cocaine use per day in past year | 4.2 ± 4.7 |
| Dollars spent per day in past year | 138.8 ± 67.4 |
| Duration of cocaine abstinence (days) | 24.3 ± 24.9 |
| Cocaine Withdrawal (CSSA) | 17.0 ± 13.5 |
| Cocaine Craving (CQ) | 16.5 ± 9.0 |
| Perceived Stress (PSS) | 21.1 ± 5.8 |
| Mean rating of drug minus neutral visual cues (cue craving) | 2.67 ± 2.19 |

Data presented in mean ± standard deviation
\*median (interquantile range)
BDI: Beck Depression Inventory
CSSA: Cocaine Selective Severity Assessment
PSS: Perceived Stress Scale
STAI: State Trait Anxiety Inventory
CQ: Cocaine Craving Questionnaire

**Figure 1.**
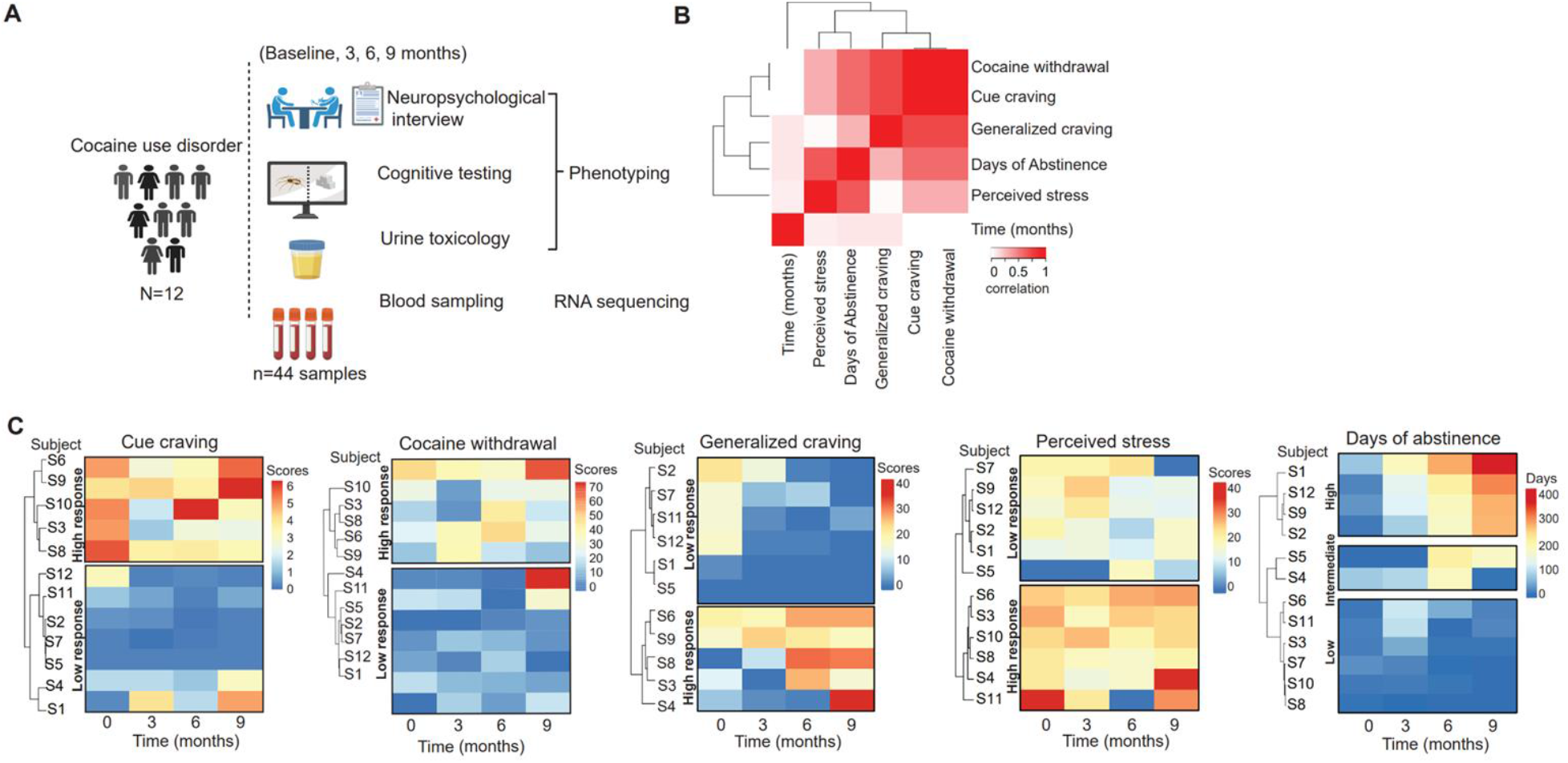
Longitudinal study design and grouping of behavioral measures. **A)**. Longitudinal study design of an outpatient cohort with CUD. Data was collected from 12 abstinence-seeking individuals with CUD consisting of drug use and biological sampling at baseline, and at 3, 6 and 9 months study visits. Negative urine toxicology for cocaine was confirmed at the baseline visit for all individuals. **B)** Canonical correlation analysis between the five selected drug use variables. **C)** Heatmaps representing pairwise correlations between the five selected variables among individuals with CUD. Hierarchical kmeans cluster analysis groups individuals into major clusters with similar scores in variables. Values of scores are shown by the color bar (top right).

To assess the relationship between these drug use variables, we performed canonical correlation analysis as shown in **Fig. 1B**. The analysis showed high collinearity between cocaine withdrawal, cue-craving and generalized craving while also revealing a moderate correlation between days of abstinence and perceived stress, and between days of abstinence, cue-craving, and cocaine withdrawal. Further, factorial analysis of these variables confirmed correlations between cue-craving, generalized craving and cocaine withdrawal as well as correlation between days of abstinence and perceived stress scores (**Fig S1**). We categorized subjects into responder groups based on their scores on the selected five drug use variables over the 4 study visit time points with k-means clustering [26]. We found two main clusters that grouped subjects into high and low responders based on scores of cue-craving, generalized craving, perceived stress and cocaine withdrawal. We also found three main clusters based on days of abstinence from cocaine use that grouped subjects to low, intermediate and high duration of abstinence as shown in **Fig. 1C**. To validate the clustering and the observed collinearity between the drug use variables, we analyzed the degree of shared respondership between the drug use variables. We observed that the subjects belonging to the cluster with high cue-craving scores were also the individuals with high cocaine withdrawal scores, demonstrating a 100% shared respondership (**Table S1, Fig. 1C**). Furthermore, subjects belonging to the cluster with high scores of generalized craving displayed an 80% shared respondership with subjects with high scores in cue craving. Lastly, subjects in the cluster with the low days of abstinence had 66.7% of shared respondership with high scores of cue-craving, and subjects with high perceived stress had 66.7% of shared respondership with high scores of cue-craving (**Fig. 1C**).

### Differential gene expression in responders grouped by drug use variables

After categorizing subjects into responder groups by scores of drug use variables, we investigated whether their peripheral blood also showed differentially expressed genes. We first corrected for the heterogeneity in gene expression caused by blood cell types using cell deconvolution analysis with CIBERSORT [27] and using the LM22 immune cell reference panel containing 22 distinct cell types (see **methods**). Deconvolution analysis revealed peripheral monocytes, natural killer resting cells, CD8 T-cells, memory CD4 resting T-cells and naive B-cells as the top five cell types contributing to a total of 0.9599 ± 0.0330 fraction of the cell-specific gene expression, while the remaining cell types showed a summated proportion of 0.0401± 0.0330 (**Fig. S2**). The monocyte population had the highest proportion (0.3150± 0.1057) for cell-specific gene expression amongst the top five cell types across subjects (**Fig. 2A**). For this reason, we included monocytes as a covariate for all downstream gene expression analyses. After rigorous QC steps (see **methods**), we also identified other technical and biological covariates such as age, gender, subject, RNA integrity (RIN) that contributed to variability in gene expression (**Fig. 2B**). Our final optimized linear mixed model was: expression ~ responder groups of drug use variables*study visit time + covariates (monocytes, age, gender, subject, RIN). Differential gene expression analysis of each variable implemented in *dream* software [28] showed no significant differences in gene expression between the responder groups of cue-craving, cocaine withdrawal, generalized craving and perceived stress scores that were time dependent at p-FDR <0.05, likely due to low sample size.

**Figure 2.**
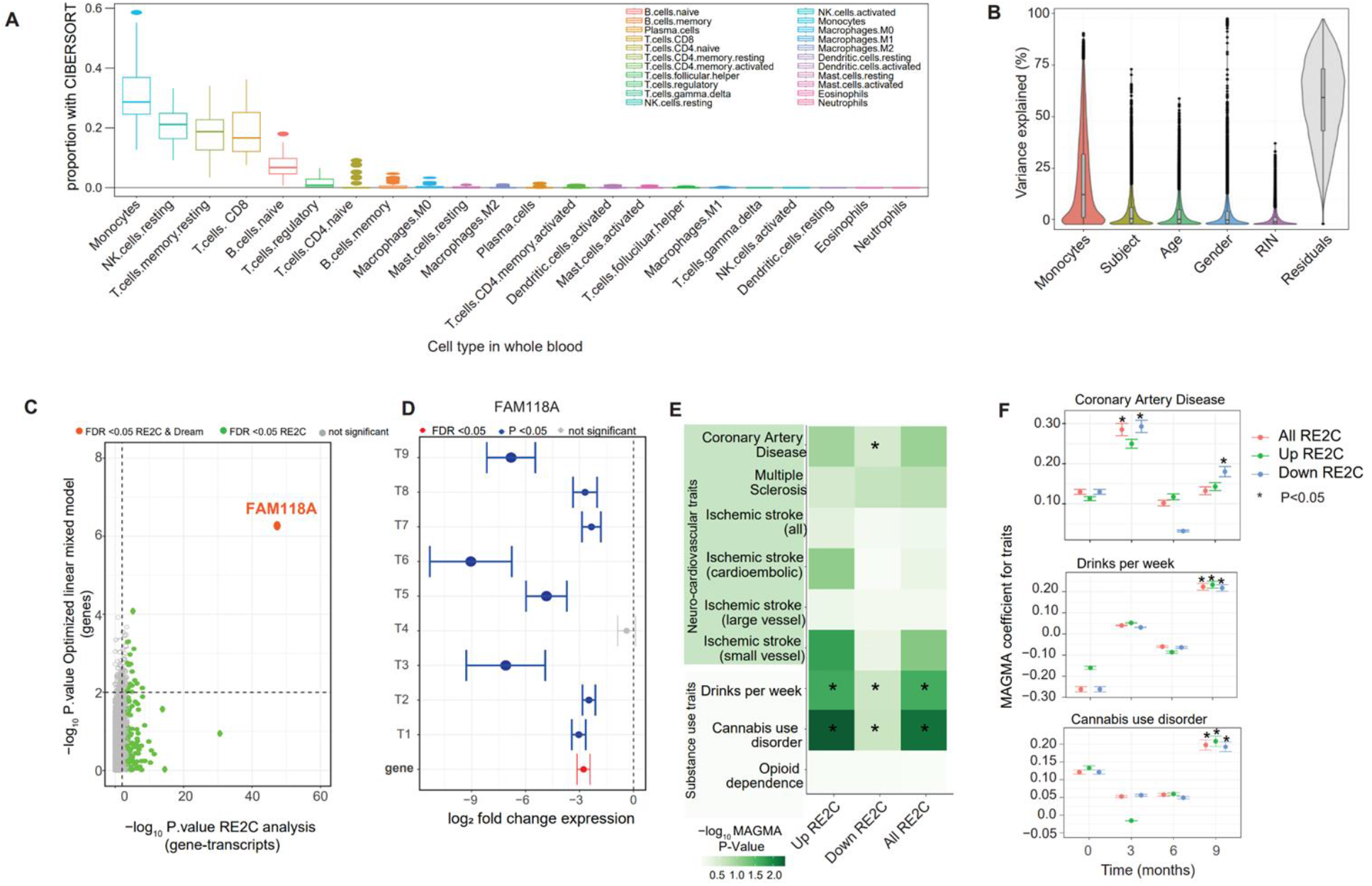
**A)**. Cell deconvolution analysis conducted with CIBERSORT demonstrates the cell types identified from whole blood, and box plots showing the cell type proportions in the study samples. The greatest gene expression variability and highest proportion is in monocytes. **B)**.Variance partition plots demonstrate additional sources of gene expression variability that are covariates for the generalized linear model. **C-D)**. Plots of high vs. intermediate days of abstinence from multivariate analysis with RE2C and concordance analysis of significant gene-transcript (i.e., transcripts T1-T9) associations at 9 months demonstrate dysregulation of FAM118A expression. **E-F)**. GWAS enrichment (MAGMA) analysis of significant transcripts shows genetic predisposition toward some substance use traits (“drinks per week” and cannabis use disorder) and coronary artery disease. Significance determined based on nominal P-value and FDR<0.05 thresholds.

### Dysregulation of transcripts in responders grouped by days of abstinence

Next, we investigated the differential expression of transcripts using our optimized linear mixed model as previously described. The analysis identified the differentially expressed gene FAM118A for high vs. intermediate days of abstinence at 9 months (**Fig. 2C**). Of note, responder groups of other drug use variables had differentially expressed transcripts although their genes were not differentially expressed (**Fig. S3-5**). These included high vs. low days of abstinence at 3 months showing differentially expressed transcript for BCORP1 (**Fig. S3**), high vs. low scores of perceived stress at 6 months for CCZ1B and SFR1 (**Fig. S4**), and high vs. low cocaine withdrawal at 9 months for XIST (**Fig. S5**). Due to a small number of samples and female subjects, the role of XIST is currently unclear.

### Gene-transcripts associations in responders grouped by days of abstinence

We next identified differentially expressed genes with their corresponding differentially expressed transcripts that we termed gene-transcripts associations. We found significant gene-transcript associations by conducting the multivariate meta-analysis of transcripts at the gene level with RE2C [29]. **Figure 2C** demonstrates an example comparing the P values from RE2C analysis and P values from differential gene expression analysis, testing for significant gene-transcript associations between high vs. intermediate days of abstinence at 9 months. As shown in **Fig 2C**, there were 116 unique differentially expressed genes that also showed significant differential expression of their corresponding transcripts (1408) at FDR < 0.05. Similarly, 181 unique genes were found at FDR < 0.05 that were linked with differences in expression of 1845 transcripts among the subjects from high vs. low abstinence days at 9 months (see **Table S2**). Notably, the differentially expressed gene FAM118A and its transcripts had significant association and concordant expression (**Fig. 2D**).

### Gene-transcript associations by days of abstinence show predisposition to substance use traits and cardiovascular disease

Enrichment analysis was conducted between significant gene-transcript associations and risk genes for clinical conditions derived from prior GWAS studies (**Fig. 2E-F, Fig. S7-S8**). Notably, risk genes in the “cannabis use disorder” and “drinks per week” GWAS traits exhibited significant enrichment for gene-transcript associations found for high and intermediate days of abstinence at 9 months (**Fig. 2E-F)**. Enrichment of risk genes in the “coronary artery disease” GWAS trait was observed for gene-transcript associations found with high vs. intermediate days of abstinence at 9 months (**Fig. 2E-F**). Moreover, significant enrichment for multiple sclerosis and small vessel ischemic stroke traits was found between intermediate vs. low days of abstinence at baseline, and coronary artery disease trait was enriched between high vs. low days of abstinence at 3 months (**Fig. S7-S8**).

### Time dependent gene co-expression networks differentiate responder groups and implicate immune regulatory mechanisms in CUD

We identified co-expressed genes that showed expression dynamics for responder groups of each drug use variable over time. To do this, we implemented weighted gene co-expression network analysis (WGCNA) [30] (**Fig. 3A**) to identify co-expressed genes labeled into modules, which was followed by non-linear regression analysis to identify modules that differentiated responder groups across time for each drug use variable. From WGCNA analysis, we identified 50 gene co-expression networks (modules), out of which, 29 modules showed biological significance with functional enrichment analysis (**Table S3**). Using non-linear regression analysis with *dream* software, we identified 8 modules that differentiated responder groups for all drug use variables at FDR<0.05. Our model used time as a non-linear variable (Gene expression ~ responder groups from clustering of drug use variables*spline(study visit time, degree=3) + identified covariates). Interestingly, genes in the tan2 module (total of 31 genes) showed expression dynamics that differentiated responder groups of days of abstinence (**Fig. 3B**). Additionally, we observed expression dynamics for responder groups with high vs low scores of cue craving and cocaine withdrawal in the firebrick and seashell4 modules (**Table S3)**.

**Figure 3.**
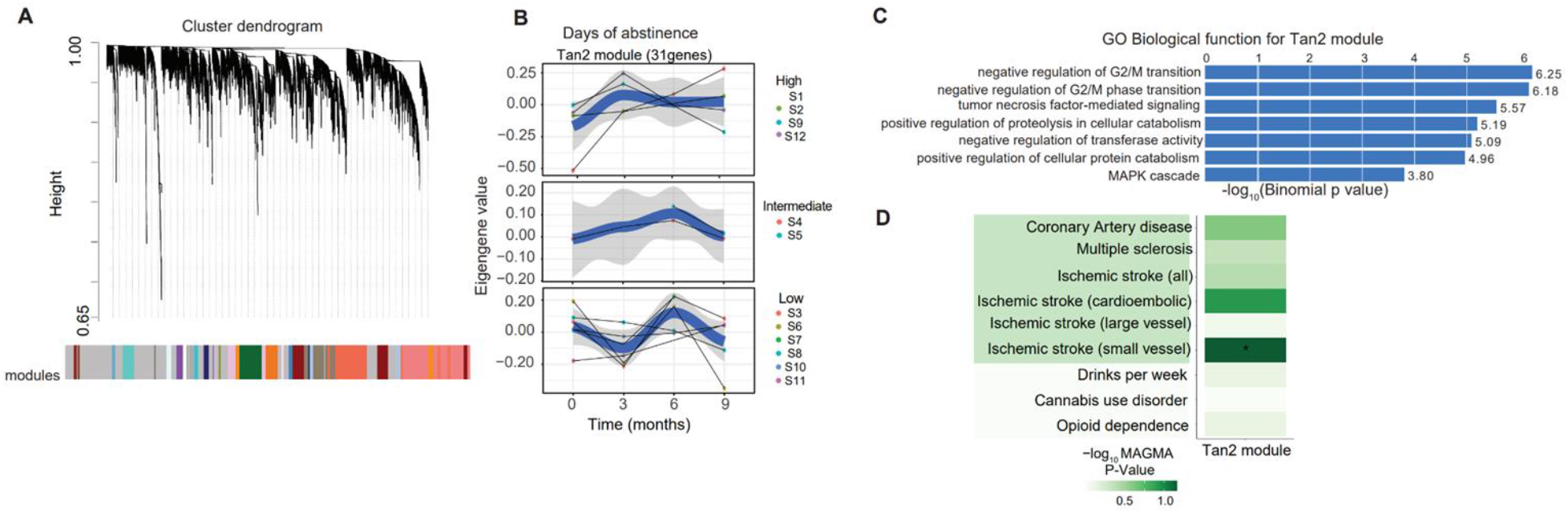
Weighted gene-co-expression (WGCNA) analysis, time series plots of module eigengenes by drug use variables with its functional significance and genetic predispositions. **A)** WGCNA network construction and organization of highly co-expressed genes in a cluster dendrogram. The genes are assigned into modules by color with eigengene values. **B)** Time series plots of the “Tan2” module showing significant changes in module eigengenes with responder groups for days of abstinence. **C)** GWAS enrichment (MAGMA) analysis showing GWAS risk loci and genetic predisposition toward ischemic stroke. Significance was determined using the nominal P-value and FDR threshold <0.05.

Co-expressed genes assigned to the tan2 module showed enrichment for functional pathways related to immune defense processes, cell cycle, DNA repair, RNA synthesis, and signal transduction (**Fig. 3C**). The tan2 module genes also showed enrichment for GWAS risk loci for small vessel ischemic stroke, a known risk factor associated with cocaine use (**Fig. 3D**) [5], of which the gene TAOK3 is a member. Similar to the tan2 module, we observed functional enrichment for immune processes, cell cycle, DNA repair, RNA synthesis, and signal transduction pathways for the firebrick module (**Fig. S10, Table S3**). The seashell4 module also showed functional enrichment for immune function and cytolysis, and enrichment for GWAS risk loci for multiple sclerosis (**Fig. S11, Table S3)**.

## Discussion

We demonstrate a streamlined analytical pipeline of a longitudinal study with bulk RNA sequencing in individuals with CUD, which can be scaled up to larger cohorts. This study is unique in that it has 4 time points of a 9-month long study in CUD. Longitudinal study designs in CUD can better identify and phenotype individuals during attempts at addiction recovery, a long and fluctuating process, and characterize mechanisms that are diagnostic or prognostic of successful abstinence from cocaine use. This study design can propel the identification of biomarkers readily accessible from whole blood in order to develop pharmacological treatments based on molecular markers that show dynamic change in tandem with abstinence from cocaine use.

With differential gene expression, we identified dysregulation of the FAM118A gene in parallel with low, intermediate, and high number of days of abstinence. While FAM118A is a lesser-known protein-coding gene, FAM118B plays an important role in Cajal body formation in cells, a specialized dynamic compartment in the nucleus involved in the formation of small nuclear ribonucleoproteins (snRNPs), pre-mRNA splicing capacity and inhibits cell proliferation [31]. FAM118B is currently studied as a cancer gene and was discovered in a GWAS of individuals with bipolar disorder and addiction [32]. We also identified gene co-expression networks which differentiated individuals with CUD over time with novel gene co-expression networks of potential functional significance associated with craving, perceived stress, cocaine withdrawal, and duration of abstinence. We found a time-dependent change in the expression of the TAOK3 gene by days of abstinence. TAOK3 belongs to the TAO (“thousand and one”) STE-20 family of serine-threonine protein kinases that are important in regulating cell stress responses, tissue homeostasis, immunity, and neurodevelopment [33]. It is found at high abundance in leukocytes and lymphoid organs, and functions as a negative regulator of inflammation and pro-inflammatory responses in macrophages [34] supporting its crucial role in immunity [35, 36]. TAOK3 also has disease relevance in neuropsychiatric disorders and substance use disorders. It is differentially expressed in post-mortem human hippocampi with CUD [37], is under epigenetic control linked to an intergenic methylated non-CpG site, and has a gene-drug interaction with opioid use disorder [38]. Therefore, our study suggests time-dependent epigenetic changes related to immune regulation, cell cycle, RNA synthesis and DNA repair during the abstinence period from cocaine use.

The relationship between transcript-gene expression revealed a genetic predisposition to relevant substance use disorders (cannabis use, alcohol use) and cardio-neurovascular conditions, all known to be comorbid in CUD [5, 39], and an ill-characterized connection to multiple sclerosis. From a biological perspective, this may reflect neuroimmune dysregulation during the course of cocaine use and abstinence [40–42], and white matter maladaptations. Neuroinflammation drives the pathogenesis of neurodegenerative diseases, including multiple sclerosis [43, 44]. Oligodendrocytes, the supportive cells that produce myelin in the white matter of the brain, are particularly vulnerable to neuroinflammation due to exposure to inflammatory cytokines such as TNF alpha and/or cell-mediated mechanisms [45] compared to other cell types, which can lead to compromised white matter integrity [46, 47]. Of note, there are several studies documenting white matter impairments using brain-imaging modalities in individuals with CUD [41, 48–50] in support of the role of neuroimmunity in CUD.

Strengths of this study include demonstrating the feasibility of a comprehensive longitudinal study characterizing the course of attempted abstinence in CUD. With our analytical pipeline, we are able to address questions regarding changes in gene expression over time differentiating complex behavioral changes during attempted abstinence. Cocaine cue-craving induces compulsion to use cocaine [11, 13] and predicts relapse to drug use. Furthermore, our findings that subjects with high scores of generalized craving display an 80% overlap with subjects with high scores in cue craving is consistent with previous findings showing that generalized craving does not fully capture drug-related craving (6,12). We also show that whole genome-based gene co-expression for cue-induced craving is also highly correlated with that for cocaine withdrawal. Future studies can determine whether examining gene expression changes during abstinence may be useful for risk stratification for remission of CUD, and identify vulnerable patients at risk for impending relapse.

Our study also demonstrates the prognostic potential of analyzing genome-wide gene expression data from whole blood. Blood is the most readily accessible sample in humans with high immune cell content, which may contain critical information that can be used for diagnostic-, prognostic- and mechanistic understanding of a given disease or perturbation [51–54]. Unlike most gene expression studies that typically rely on analysis of post-mortem brain tissues, longitudinal transcriptomics can show dynamic behavioral changes in living subjects, with gene expression changes in whole blood. We applied gene co-expression network analysis as a way to demonstrate time-dependent dysregulations in individuals with CUD. We also applied differential gene-transcript associations as a strategy to identify genetic predispositions to comorbid clinical conditions with CUD that may prove useful to improve the treatment of patients during abstinence. These results highlight the importance of screening for cardio-neurovascular diseases in individuals currently in treatment for CUD to prevent future morbidity and mortality.

This study is preliminary in that the sample size of subjects is limited along with a small sample size of gene expression data from whole blood. Although analysis of whole blood may not directly reflect perturbations within the central nervous system, there is increasing evidence of the utility of these blood markers in the pathway from discovery to pharmacological treatments of addiction [55–57]. The study cohort is homogenous predominantly of older (over 45 years) and of African descent. Although it reflects the geographic location the study was conducted and treatment-seeking individuals, it may not reflect the demographics of the general cohort of individuals with CUD [58]. Indeed, expansion of the sample size will help towards generalizability by increasing the number of subjects and growing the sample cohort, inclusion of a control group without CUD, and incorporating populations from other geographic locations.

In conclusion, it is currently very challenging to predict the conditions that either promote abstinence or predict relapse to cocaine use, which reflects the highly phenotypic heterogeneity in CUD. A longitudinal study design best captures the degree of phenotypic heterogeneity, especially during the abstinence period, however it can be very complex to conduct. Our feasible analytical pipeline for complex longitudinal studies with neuropsychological phenotyping and blood-based transcriptomics, demonstrated time-dependent dysregulations in gene-transcription expression and gene expression of molecular and clinical significance. In the future, the combinatorial use of molecular, behavioral and neuropsychological biomarkers may improve methodologies and treatment modalities used in CUD [24] by predicting either successful or poor treatment responses, clinical comorbidities, and vulnerable time windows for relapse to cocaine use.

## Acknowledgements

We would like to thank Gabriel Hoffman for his support with computational analyses. All figures were assisted with the use of BioRender.com. This work was supported in part through the computational and data resources and staff expertise provided by Scientific Computing and Data at the Icahn School of Medicine at Mount Sinai and supported by the Clinical and Translational Science Awards (CTSA) grant UL1TR004419 from the National Center for Advancing Translational Sciences. Research reported in this publication was also supported by the Office of Research Infrastructure of the National Institutes of Health under award number S10OD026880 and S10OD030463.

## Author Contributions

Drs. Nwaneshiudu and Girdhar had full access to all the data in the study and take responsibility for the integrity of the data and the accuracy of the data analysis. Study concept and design: All authors.

Acquisition, analysis, or interpretation of data: All authors.

Drafting of the manuscript: All authors

Critical revision of the manuscript for important intellectual content: All authors.

Statistical analysis: Nwaneshiudu, Girdhar.

Obtaining funding: Roussos, Goldstein.

Administrative, technical, or material support: Roussos, Goldstein.

Study supervision: Roussos, Alia-Klein, Goldstein.

## Data and Code availability

all publicly available softwares are noted in the Methods session. Data will be available upon request to study investigators.

## Supplementary Material

[Supplementary data and methods](https://docs.google.com/document/d/1pYzLQeL3VBfdJfIRbqI2LhXHi1l2z_8wHy31HfqK9KQ/edit)

