## Supplementary data and methods for "Dynamic alterations in gene co-expression networks and gene-transcript associations characterize co-morbidities in cocaine use disorder"

This PDF file includes:

- Materials and methods
- Supplementary Figs. 1-10
- Supplementary Tables S1-S3

**Materials and methods**

*Subjects*

Protocols within the study were approved by the institutional review board of the Icahn School of Medicine at Mount Sinai, and were conducted in accordance with standards presented in the 1964 Declaration of Helsinki. Twelve subjects with CUD (not in treatment) provided written, informed consent for participation in accordance with the local institutional review board. A comprehensive diagnostic interview based on DSM5 criteria was conducted to determine history of CUD, psychological and medical histories and a detailed description of the study protocol and methods has been published [[1, 2]](https://paperpile.com/c/R5hdBS/cAX0+1mQL). Briefly, participant samples and assessments were collected between November, 2020 and July, 2021 undergoing comprehensive phenotyping with neuropsychological assessments, urine toxicology, cognitive testing and blood sample collection during their baseline visit and repeatedly at 3, 6, and 9 months thereafter. The baseline visit was when all participants had negative urine drug screens for cocaine, and were abstinent from recent cocaine use for >3 days. All aspects of the study protocol, as well as the present analysis, were approved by the local Institutional Review Board, and all patients provided written, informed consent. Data analysis for the current study was performed from March 2022 to October 2023.

*Assessments*

Clinical Psychological (CP) battery: All tests were presented in the same fixed order for all subjects. The complete battery was administered in a quiet room, with all subjects completing the CP battery in one sitting. To minimize fatigue, subjects were encouraged to take a short break toward the middle of the battery. Individuals that use cigarettes were asked to refrain from smoking throughout the CP session and the short break. The following cocaine craving measures were collected; generalized cocaine craving [[3]](https://paperpile.com/c/R5hdBS/xk0w), and cocaine craving in response to visual cues (drug, neutral, pleasant) [[4]](https://paperpile.com/c/R5hdBS/H90F). Additional measures of self-reported days of abstinence from cocaine use, symptoms of cocaine withdrawal and perceived stress scales were also collected using the cocaine selective severity assessment scale[[5]](https://paperpile.com/c/R5hdBS/e8Dc), and perceived stress scale (PSS-10) [[6, 7]](https://paperpile.com/c/R5hdBS/ri1g+0v07) respectively.

*Sample collection and processing*

Gene expression data from whole blood [[8, 9]](https://paperpile.com/c/R5hdBS/ZrUw+xnb4): Peripheral blood samples (N=48 samples overall; i.e., n=12 individuals at 4 timepoints) collected from subjects during their baseline, 3, 6, 9 months visits to generate cDNA libraries for sequencing. Total RNA was extracted, isolated and quantified from the blood samples. The cDNA library was synthesized using random hexamers, ligated with appropriate adaptors to barcode the samples for high throughput sequencing. The cDNA libraries were sequenced on the Illumina HiSeq 2500 System. The raw sequence reads were aligned to human genome database hg19 with STAR aligner (v.2.7.3a) and featureCounts16 were used to quantify the gene expression at the gene level based on Ensembl gene model GRCh37.70.  We then quantified both gene- and exon-level expression in the cDNA library. The gene level read counts data was normalized as counts per million (CPM) using the trimmed mean of M-values normalization (TMM) method with voom from the limma R package [[10]](https://paperpile.com/c/R5hdBS/9Ur2), to adjust for sequencing library size differences. Sequencing quality control (QC) metrics were calculated by fastqc (v0.11.8) and Picard Tools (v2.22.3) [[11]](https://paperpile.com/c/R5hdBS/z0Ui). The raw reads were filtered after the above quality control assessments, to remove low quality variant sites using recommended parameters. Also, low-expression genes and transcripts were removed, keeping genes and transcripts with counts per million >1 in 10% of the samples. Further genome-wide quality control analysis was conducted using multidimensional scaling (MDS) [[12]](https://paperpile.com/c/R5hdBS/HrGD) and principal component (PCA) [[13]](https://paperpile.com/c/R5hdBS/1xwM) analyses which reduced overall samples for further analysis to N=44.

### *Analysis of study metadata, genes and transcripts in blood*

Grouping of study metadata drug use variables: The grouping of study metadata was conducted to facilitate harmonizing the metadata with the transcriptomic data. Cocaine craving measures (cue craving, generalized craving), perceived stress, days of abstinence and cocaine withdrawal measures as a function of time were factorized using hierarchical k-means clustering [[14]](https://paperpile.com/c/R5hdBS/zd0T) into two groups (low and high responders) as a function of time. The days of abstinence from cocaine use as a function of time was also factorized into 3 distinct groups; High days-sustained abstinence, intermediate days and low days- recent cocaine use. Exploratory factorial analysis [[15]](https://paperpile.com/c/R5hdBS/JWgB) was conducted on the five drug use variables for harmonization.

*Quality control of gene transcripts*: Gene level and genome wide QC analysis to verify sequencing was previously described [[16]](https://paperpile.com/c/R5hdBS/6VaU). Briefly, retained samples in the RNA library were assessed for their RNA Integrity Number (RIN), intergenic rate, total read pairs, intronic rate, mapped read pairs, and ribosomal RNA rates. Using PCA analysis of autosomal genes identified outliers, and distinct clusters correspond to technical differences in the RNA library preparation. After that, gene raw counts were filtered to include only those on autosomes with counts per million reads (CPM) > 0.1 in at least 25% of samples. Count-level quantifications were corrected for library size by using trimmed mean of M-values (TMM) normalization and were log_2_ transformed.

*Cell deconvolution*: Cell type proportions were calculated with CIBERSORTx [[17, 18]](https://paperpile.com/c/R5hdBS/KqQD+0N26), which uses the LM22 as a reference gene expression panel with 22 different cell types (naive B cells, memory B cells, Plasma cells, CD8	T cells, CD4 naive T cells, resting memory CD4 T cells, activated memory CD4 T cells,	follicular helper	T cells, regulatory T cells, gamma delta	T cells, resting natural killer (NK) cells, activated NK cells, Monocytes, M0 Macrophages, M1 Macrophages, M2 Macrophages, resting Dendritic cells, activated Dendritic cells, resting Mast cells, activated Mast cells, Eosinophils and Neutrophils) for computation. The cell type with the greatest gene expression variability was included as a covariate in the linear model.

*Selection of biological and technical covariates for optimized model*: We then used the multivariate information criteria (mvIC) [[19]](https://paperpile.com/c/R5hdBS/WXSU) to assess significant variables that influenced the most gene expression variability, using forward stepwise regression to further optimize the model. MVIC is an extension of the standard Akaike (AIC) or Bayesian Information Criterion (BIC) for multivariate regression modeling that analyzes high dimensional datasets with many response variables. Using forward stepwise regression, it automatically selects only the variables with significant correlations with the principal components of the data. With the final model, we computed the variance partition to assess the percentage of gene expression variability explained by these significant variables.

*Differential expression of genes:* With the dream software [[20]](https://paperpile.com/c/R5hdBS/vNOp), the optimized generalized linear mixed model was applied that accounts for fixed and random effects in the equation Gene expression ~ 1|subject + group*time, while controlling the false positive rate and allows for cross-individual hypothesis testing [[20, 21]](https://paperpile.com/c/R5hdBS/vNOp+ja3R). Residual degrees of freedom were estimated with the Satterthwaite approximation. For all gene expression analyses, we established statistical significance using a Benjamini–Hochberg false discovery rate (FDR) correction of <5% [[22]](https://paperpile.com/c/R5hdBS/x5iM). We added the drug use variables of interest with time as an interaction term to identify time-dependent differentially expressed genes.

*Differential expression of transcripts*: After filtering and normalization steps as above, genes were filtered to include only those on autosomes longer than 250 base pairs with transcripts per million reads (TPM) > 0.1 in at least 25% of samples. Count-level quantifications were corrected for library size by using trimmed mean of M-values (TMM) normalization and were log_2_ transformed. We then imputed the transcript data into the optimized linear mixed model using the dream software, adding the drug use variables with time as an interaction term to identify time-dependent differentially expressed transcripts. We then used multivariate testing (mvTest) from the dream software [[20]](https://paperpile.com/c/R5hdBS/vNOp) to examine association of differentially expressed transcripts for each differentially expressed gene. The multivariate test computes the correlations between estimated regression coefficients across multiple responses based on REC2, a powerful random effects method for meta-analysis of GWAS studies [[23]](https://paperpile.com/c/R5hdBS/QPzt) that accounts optimally for correlations, and focusing on heterogeneous effects conditioned on the application of fixed effects. Outside of GWAS, this method can be applied to a wide range of study designs such as cross-disease or cross-population studies [[23]](https://paperpile.com/c/R5hdBS/QPzt). We applied this analysis to identify the list of differentially expressed gene-transcript(s) with significant associations for each drug use phenotype. We were also able to observe the concordance of expression between the differentially expressed genes and their transcript variants.

RE2C utilized estimated effect sizes of transcripts derived from our optimized linear mixed model as described above. Unlike conventional meta-analyses, this analysis relaxes the assumption of statistical independence among estimated effect sizes of transcripts. It incorporates the inter-correlation matrix between effect sizes of the transcripts and models them as random effects using the method from Han et.al. (24) and finally provides the association statistics at gene level. Consistent with our previous objective, we sought to explore the differential expression of transcripts within identified subject clusters associated with neuropsychological variables over clinic visit times.

*Gene set enrichment analysis with MAGMA*: To test the enrichment of associated differentially expressed gene-transcript(s), we used MAGMA software [[24]](https://paperpile.com/c/R5hdBS/dym2). This gene and gene set enrichment analysis is based on multiple linear principal components regression that allows generalizable analysis of both continuous properties of genes and multiple gene sets. Individual genes are aggregated to groups of genes that share certain biological, functional or clinically relevant characteristics. We used MAGMA to assess overlap between differentially expressed transcripts and a large reference GWAS database containing genes and gene-sets known to be associated with clinical conditions and anthromorphometric features, to determine genetic predisposition to clinical conditions based on differentially expressed gene-transcript associations. An F-test and p-value was then computed for each differentially expressed gene-transcripts and its associated clinical condition(s).

*Gene-gene co-expression analysis*: Weighted gene expression correlation network analysis was performed using the WGCNA R package [[25]](https://paperpile.com/c/R5hdBS/x38L). The package provides a comprehensive set of functions (eigengene network construction, module detection with unsupervised clustering and dynamic tree cutting, formulating relationship of modules to drug use data), for performing and describing the data structures of highly correlated gene transcripts integrated with other high-dimensional data. We applied this analysis to identify modules and their component genes with significant gene expression variation over time. We then examined module-trait and time associations with generalized and cocaine- cue reactive craving, days of abstinence, cocaine withdrawal and perceived stress.

*Time spline analysis*: Using the modules of biological significance with their component genes, we used fitted spline curve modeling with Dream [[20]](https://paperpile.com/c/R5hdBS/vNOp) to test for significant gene co-expression change over time with the generalized linear model Expression ~ drug use variables responder groups*basic spline (time, degree = 3) + covariates (gender, subject, monocytes, age, RIN). We then assessed for differences in time dependent variation by groupings of generalized and cocaine- cue reactive craving, days of abstinence, cocaine withdrawal and perceived stress. We then applied functional enrichment analysis using the GREAT tool (Genomic Regions Enrichment of Annotation) to determine if the co-expressed genes showing significant time-dependent expression also had shared biological significance.

*Functional enrichment analysis*: To determine the biological function of component genes in modules, functional enrichment analysis was performed using GREAT [[26]](https://paperpile.com/c/R5hdBS/sqtp).

**Figure S1**: Factorial analysis of drug use variables in individuals with CUD.

**Figure S2**: Cell-type proportion summation from whole blood in individuals with CUD.

**Figure S3**: Differential transcript expression by greater vs. least abstinent days at 3 months and multivariate analysis with RE2C demonstrating significant gene-transcript associations.

**Figure S4**: Differential transcript expression by high vs. low perceived stress scores at 6 months and multivariate analysis with RE2C demonstrating significant gene-transcript associations.

**Figure S5**: Differential transcript expression by high vs. low scores of cocaine withdrawal at 9 months and multivariate analysis with RE2C demonstrating significant gene-transcript associations.

**Figure S6**: Concordance analysis of CCZ1B gene and its transcripts (T1-T6) expression levels between high vs. low perceived stress scores at 6 months. Overall there is significant concordant change in expression with CCZ1B gene and its transcripts.

**Figure S7**: GWAS enrichment analysis with MAGMA of significant RE2C transcripts between days of abstinence with time of clinic visit. Heatmap of P-values of clinical traits and effect size of significant clinical traits based on nominal P-value (*) thresholds demonstrate genetic predispositions toward cannabis use disorder, drinks per week, coronary artery disease, multiple sclerosis and ischemic stroke involving small vessels.

**Figure S8**: GWAS enrichment analysis with MAGMA of significant RE2C transcripts between days of abstinence with time of clinic visits. Effect size of significant clinical traits based on nominal P-value (*) thresholds demonstrate significant genetic predispositions toward cannabis use disorder, drinks per week, coronary artery disease, multiple sclerosis and ischemic stroke involving small vessels.

**Figure S9**: A). Time dependent gene co-expression module eigengene values over time with significant responder differences with cue-induced craving and cocaine withdrawal using spline analysis. B). Functional significance of firebrick module showing significant enrichment with immune processes.

**Figure S10**: Time dependent gene co-expression module eigengene values over time with significant responder differences with cue craving and cocaine withdrawal using spline analysis. Functional significance of seashell4 module showing significant enrichment with immune processes. GWAS enrichment analysis demonstrating significant genetic predisposition towards multiple sclerosis.

| **Shared responders** | | **High response (%)** | **Low response (%)** |
| --- | --- | --- | --- |
| **drug use variable 1** | **drug use variable 2** |  | |
| Cocaine withdrawal | Cue craving | 5/5 (100%) | 7/7 (100%) |
| Generalized craving | Cue craving | 4/5 (80%) | 6/6 (100%) |
| Perceived stress | Cue craving | 4/6 (66.7%) | 5/6 (83.3%) |
| Days of Abstinence (high) | Cue craving | 1/4 (25%) | 3/4 (75%) |
| Days of Abstinence (intermediate) | Cue craving | 0/2 (0%) | 2/2 (100%) |
| Days of Abstinence (low) | Cue craving | 4/6 (66.7%) | 2/6 (33.3%) |
| Cocaine withdrawal | Generalized craving | 4/5 (80%) | 6/6 (100%) |
| Cue craving | Generalized craving | 4/5 (80%) | 6/6 (100%) |
| Perceived stress | Generalized craving | 4/6 (66.7%) | 5/6 (83.3%) |
| Days of Abstinence (high) | Generalized craving | 1/4 (25%) | 3/4 (75%) |
| Days of Abstinence (intermediate) | Generalized craving | 0/2 (0%) | 2/2 (100%) |
| Days of Abstinence (low) | Generalized craving | 3/6 (50%) | 3/6 (50%) |

**Table S1**: Percentage of individuals in shared responder groups to drug-induced cue craving and generalized craving by drug use variable. Calculated based on number of matched individuals with response in drug use variable 1 and 2 divided by number of individuals with response in drug use variable 1

**Table S2**: Differential transcript expression and multivariate analysis with RE2C by shared responder groups of drug use variables. Number of significant RE2C gene-transcript associations used for imputation for GWAS enrichment analysis is also demonstrated.

| **Time dependent WGCNA module** | **Module with functional significance** | **Functional pathway** | **drug use variable** | **Number of time dependent genes** |
| --- | --- | --- | --- | --- |
| firebrick | Functionally enriched | Immune regulation | Cocaine withdrawal/Cue craving  Cue craving | 23  23 |
| tan2 | Functionally enriched | Cell cycle, signal transduction | Cocaine withdrawal  Generalized craving  Cue-craving  Days of abstinence | 2  5  2  10 |
| seashell4 | Functionally enriched | Immune response | Cue craving  Days of abstinence  Cocaine withdrawal | 45  2  45 |
| cornsilk | Not functionally enriched | - | Cue craving  Cocaine withdrawal | 2  2 |
| darkseagreen | Functionally enriched | ER stress, protein folding | Perceived stress  Cue craving  Cocaine withdrawal | 1  3  3 |
| oldlace | Not functionally enriched | - | Perceived stress  Generalized craving | 3  2 |
| darkturquoise | Functionally enriched | Cell cycle regulation, brain/skull development | Perceived stress | 1 |
| royalblue3 | Not functionally enriched | - | Days of abstinence | 3 |
| black | Functionally enriched | Immune defense mechanisms | Days of abstinence | 1 |
| coral4 | Functionally enriched | RNA synthesis | Days of abstinence | 2 |

**Table S3**: Gene co-expression analysis with WGCNA and spline analysis with dream demonstrate time-dependent modules and genes that differentiate responders by drug use variables. Time-dependent modules with functional significance are also demonstrated.

11.	Institute B. \textquotedblleftPicard Toolkit\textquotedblright, Broad institute, GitHub repository. Picard Toolkit. 2019. 2019.

12.	Davison ML, Sireci SG. 12 - Multidimensional Scaling. In: Tinsley HEA, Brown SD, editors. Handbook of Applied Multivariate Statistics and Mathematical Modeling, San Diego: Academic Press; 2000. p. 323–352.

13.	Stein SAM, Loccisano AE, Firestine SM, Evanseck JD. Chapter 13 Principal Components Analysis: A Review of its Application on Molecular Dynamics Data. In: Spellmeyer DC, editor. Annual Reports in Computational Chemistry, vol. 2, Elsevier; 2006. p. 233–261.

14.	Pérez-Ortega J, Almanza-Ortega NN, Vega-Villalobos A, Pazos-Rangel R, Zavala-Díaz C, Martínez-Rebollar A. The K-means algorithm evolution. Introduction to Data Science and Machine Learning. 2019. 3 April 2019. https://doi.org/[10.5772/INTECHOPEN.85447](http://dx.doi.org/10.5772/INTECHOPEN.85447).

15.	Fabrigar LR, Wegener DT. Exploratory Factor Analysis. OUP USA; 2012.

16.	Hoffman GE, Bendl J, Voloudakis G, Montgomery KS, Sloofman L, Wang Y-C, et al. CommonMind Consortium provides transcriptomic and epigenomic data for Schizophrenia and Bipolar Disorder. Scientific Data. 2019;6:180.

17.	Chen B, Khodadoust MS, Liu CL, Newman AM, Alizadeh AA. Profiling Tumor Infiltrating Immune Cells with CIBERSORT. Methods Mol Biol. 2018;1711:243–259.

18.	Newman AM, Steen CB, Liu CL, Gentles AJ, Chaudhuri AA, Scherer F, et al. Determining cell type abundance and expression from bulk tissues with digital cytometry. Nat Biotechnol. 2019;37:773–782.

19.	Hoffman G. mvIC: Multivariate Information Criteria to identify important predictors in high dimensional multivariateregression. R package version 1.6.3. mvIC: Multivariate Information Criteria to Identify Important Predictors in High Dimensional Multivariateregression R Package Version 1 6 3. 2022.
